# Metabolic redundancy is required for microbial polyethylene assimilation

**DOI:** 10.64898/2025.12.10.693370

**Authors:** Theo Obrador-Viel, Rocío D. I. Molina, Robyn J. Wright, Maria del Mar Aguiló-Ferretjans, Vinko Zadjelovic, Balbina Nogales, Rafael Bosch, Joseph A. Christie-Oleza

## Abstract

Polyethylene (PE) is amongst the most recalcitrant synthetic polymers, and only a limited number of microbes have been shown to utilise it as their sole carbon and energy source. Here, we investigated the metabolic basis enabling the efficient assimilation of PE oxidised scission products and its prevalence in microbial communities naturally colonising plastic surfaces. Metabolomic profiling of weathered PE (W-PE) leachates revealed a highly diverse pool of oxidised aliphatic compounds varying in chain length and oxidation state. Different plastic-degrading bacteria consumed this complex mix of metabolites to distinct extents, with consumption efficiency positively correlating with the number of redundant genes associated with the β-oxidation pathway in their genomes. Comparative proteomic analysis of two *Alcanivorax* species exhibiting contrasting PE-leachate consumption capabilities confirmed that this functional redundancy was fully activated in response to the chemically complex PE-derived substrate pool. In contrast, it remained largely uninduced in the presence of the single, structurally simple alkane hexadecane. Hence, our results indicate that efficient PE assimilation requires a broad and redundant enzymatic repertoire capable of funnelling structurally diverse oxidised aliphatic intermediates through β-oxidation. Metagenomic analysis of plastisphere communities further revealed enrichment of fatty acid degradation genes in biofilms colonising both pristine and weathered PE—as expected more strongly in W-PE—compared with wood and surrounding water controls, supporting the ecological relevance of this mechanism for PE biodegradation. Together, these findings identify β-oxidation metabolic redundancy as a key trait underpinning microbial PE assimilation and suggest that plastic degradation may be occurring under natural environmental conditions.

## INTRODUCTION

Single-use plastics have become an irreplaceable part of modern society due to their cost-effectiveness and convenience. However, improper waste management has resulted in significant social and environmental challenges. Since plastic debris was first detected on the ocean’s surface in the early 1970s (Carpenter and Smith, 1972), marine plastic pollution has emerged as an escalating concern, posing a global threat to already fragile ecosystems. Despite increased public awareness and recent political initiatives, plastic waste entering natural ecosystems is expected to continue rising (Williams and Rangel-Buitrago, 2022; García Rellán *et al*., 2023).

Polyethylene (PE) is one of the most abundant polymers in the environment due to its high production –over 26 % of global plastic production– and its low density (0.88-0.96 g cm^-3^) which increases its dispersal in waterbodies (Wright *et al.,* 2020). Its simple linear structure of carbon– carbon chains occur in two primary forms: low-density (LD-PE), with branched chains that confer flexibility for packaging and bags, and high-density (HD-PE), with a more crystalline, rigid structure suited for durable products (Ghatge *et al*., 2020). Owing to its ubiquity and persistence, increasing attention has focused on microbial species capable of colonising and degrading PE.

PE shares structural similarities with linear alkanes, suggesting *a priori* that bacterial alkane degradation pathways may also act on PE. Although both are chemically inert, many microorganisms can metabolise alkanes as an energy-rich carbon source. Alkanes degradation typically begins with a hydroxylation to alcohols, which are subsequently oxidised to aldehydes and fatty acids, and ultimately processed via the β-oxidation pathway (Rojo, 2009). An alternative subterminal pathway involves ketone formation and conversion to esters via a Baeyer-Villiger monooxygenase, followed by hydrolysis into carboxylic acids and alcohols (Ji *et al*., 2013).

Despite structural parallelisms, PE and alkanes differ profoundly. The longest metabolised alkanes reported reach C36 (Wang and Shao, 2014; Xiang *et al*., 2022), whereas PE polymer chains extend to thousands of carbons, which are excessively long to fit into monooxygenase active sites or access inner-membrane enzymes (Guo *et al*., 2023). Furthermore, PE’s high hydrophobicity and crystallinity further limit microbial access and biodegradation (Chow *et al*., 2023). PE, however, undergoes abiotic (and biotic) weathering in the environment, leading to random oxidation and chain scission that generate shorter oxidised aliphatic fragments (Gewert *et al*., 2018) which specialised microbes can then assimilate (Romera-Castillo *et al.,* 2018).

Alkane-degrading bacteria –or so-called obligate hydrocarbonoclastic bacteria (OHCB)– are widespread in the environment and often bloom in response to oil pollution. Well-known alkane degraders include the marine genera *Alcanivorax*, *Marinobacter* and *Halomonas* (Wang *et al*., 2025; Yakimov *et al*., 2007) as well as the terrestrial genera *Pseudomonas* and *Rhodococcus* (Gunasekera *et al*., 2025; Kok et al., 1989; Sun *et al*., 2025). Members of these and other alkane-degrading taxa are frequently detected in the plastisphere, i.e. biofilms that develop on plastics. For instance, *Oleiphilus*, *Roseobacter*, and *Aesturiibacter* were enriched in weathered PE (W-PE) biofilms after short incubations in seacoast environments (Erni-Cassola *et al*. 2020), while *Alcanivorax*, *Marinobacter*, and *Arenibacter* were found on LD-PE (Delacuvellerie *et al*., 2019); putative hydrocarbon degraders comprised up to 17.4% of a PE biofilm community (Vaksmaa *et al*., 2021). Such findings often fuel speculation on plastic biodegradation in the environment.

Despite the increasing number of studies on PE degradation, only a few have characterised its biological mechanisms. The terrestrial strain *Rhodococcus rhodochrous* ATCC29672 was shown to assimilate oxidised PE oligomers (Eyheraguibel *et al*., 2017) and even mineralise ¹³C-labelled PE (Goudriaan *et al*., 2023), with alkane-degradation and β-oxidation pathways upregulated during the process (Gravouil *et al*., 2017). Similarly, a proteomic analysis of *Alcanivorax* sp. 24 revealed the induction of an extensive array of fatty acid degradation enzymes when grown on W-PE, indicating the uptake of leached molecules, their oxidation to carboxylic acids, and subsequent processing via CoA activation and β-oxidation to acetyl-CoA, supplemented by auxiliary enzymes that channelled more complex intermediates into this pathway (Zadjelovic *et al*., 2022).

Nevertheless, what defines an efficient PE biodegrader and which metabolic routes enable its assimilation remain open questions in plastisphere research. In this study, we characterised the bioavailable byproducts released from W-PE to determine the substrates accessible to microbes and the metabolic pathways required for their utilisation. We then linked the growth of a collection of marine bacterial isolates on W-PE with their genomic potential to identify the metabolic traits associated with effective PE assimilation. Finally, we assessed whether these metabolic pathways were enriched in PE plastisphere metagenomes, underscoring their ecological relevance and use in future plastic biodegradation research.

## MATERIALS AND METHODS

### Polymers used and material preparation

Experiments were performed using LD-PE pellets (approx.. 4 mm, Sigma-Aldrich®) with a molecular weight of 122,900 g/mol (range: 100,000-250,000) as specified by the manufacturer. Polymer was weathered by incubating the LD-PE pellets in glass flasks covered with foil at 80 °C in a dry oven for 6 months.

### Attenuated Total Reflectance Fourier-Transform InfraRed (ATR-FTIR) spectroscopy

The polymer oxidation state was analysed by Attenuated Total Reflectance Fourier-Transform InfraRed (ATR-FTIR) spectroscopy. Spectra were obtained using an FTIR spectrometer (Tensor 27, Bruker) in transmission mode at the wave-number range between 400-4,000 cm^−1^. The number of scans was 32, with 8 seconds *per* scan, with a resolution of 4.0 cm^−1^ and an interval of 1.0 cm^−1^. Spectra were smoothed and baseline-corrected using the online tool in the OpenSpecy web version (Cowger *et al*., 2021). Then, the weathering intensity was evaluated by calculating the carbonyl index corresponding to the widely used carbonyl peak at 1712 cm^−1^, normalised to the intensity of the control region 1450 cm^−1^, which is not affected by oxidation (Almond *et al*., 2020).

### Leachate acquisition and sample collection

Pristine and weathered plastic pellets were fragmented with a scalpel into approx. 1 mm pieces, weighed, disinfected with ethanol and dried over 3 days at 30 °C. The resulting fragments were transferred to acid-washed (10 % HNO_3_ for 1 h and thoroughly rinsed with distilled water) and autoclave-sterilised 100 mL Erlenmeyer flasks, after which 30 mL of sterile Milli-Q water was added to obtain a final plastic concentration of 0.3 % w/v. Water blanks were processed in parallel for downstream baseline determination. To simulate the leaching of compounds from plastic and their availability to bacteria over time, the flasks were incubated at 30 °C with orbital shaking at 170 rpm for 14 days. At defined timepoints (0, 20 min, 1 h, 6 h, 1 d, 3 d, 7 d and 14 d), 1.5 mL aliquots of the supernatant were collected, centrifuged for 3 min at 16,200 ×g to remove residual plastic fragments, and stored at −20 °C until further analysis: 1 mL for total dissolved organic carbon determination and 200 µL for metabolomic analysis as described below. Flasks were kept sterile until the end of the experiment.

### Total dissolved organic carbon (DOC) measurements

The organic carbon leached by the plastics was measured with a TOC-C CPH analyser (Shimadzu) equipped with an ASI-L 24 MLS autosampler. 1 mL samples were added to 9 mL of Milli-Q water in acid-washed vials and analysed in non-purgeable organic carbon (NPOC) mode with 1.5 % hydrochloric acid (HCl) addition; 50 μL of injection volume and three technical replicates with 2 washes between samples. Sparge gas flow was 80 mL min^−1,^ with a sparge time of 1.5 min. The exact concentration was calculated with a calibration curve prepared from a 1,000 ppm terephthalate solution.

### Metabolomic analysis by UHPLC-MS/MS

Molecular formulae of compounds leached from pristine and W-PE were determined using ultra-high performance liquid chromatography tandem mass spectrometry (UHPLC-MS/MS). For sample preparation, 200 µL aliquots were dried using a SpeedVac (Eppendorf® Concentrator Plus) at 40 °C for 30 min, then reconstituted in 20 µL of 5 % acetonitrile (Merck) containing 1 ‰ formic acid (Merck) prior to analysis. UHPLC-ESI-MS/MS analysis was performed on an UltiMate 3000 system (Thermo Fisher Scientific) coupled to a Q-Exactive Hybrid Quadrupole-Orbitrap mass spectrometer (Thermo Scientific). The injection volume was 5 μL.

Chromatographic separation was achieved using a reverse-phase C18 core-shell column (Luna Omega, 100 × 2.1 mm, 1.6 μm; Phenomenex, Torrence, CA) preceded by a C18-security guard ultra-cartridge (2.1 mm; Phenomenex). The mobile phases consisted of (A) 0.1 % (v/v) formic acid in Milli-Q water and (B) 0.1 % (v/v) formic acid in acetonitrile, delivered at 0.2 mL min^-1^. The gradient started at 5 % solution B for 5 min, increased linearly to 55 % over 5 min, held for 2.5 min, then ramped to 90 % over 3.5 min and held for 5 min. Full MS scans were acquired in both positive and negative ion modes over an *m/z* range of 85–1,275 at a resolution of 70,000. Data-dependent MS/MS scans targeted the five most intense peaks *per* cycle with a dynamic exclusion of 3 s, using an *m/z* range of 200-2,000 and a resolution of 17,500. The resulting data were analysed with the Compound Discoverer 2.0 Software (Thermo Fisher Scientific). Parameters for molecule prediction were as follows: mass tolerance of 10 ppm, intensity tolerance of 30 %, minimum peak intensity of 5×10^6^, signal-noise ratio threshold of 3, maximum elements count of C100, H202, and O50. Named annotations of the compounds were assigned based on ‘Predicted Compositions’, ‘mzCloud™’ (mzcloud.org) and ‘ChemSpider’ (Pence and Williams, 2010). Statistical analysis of the MS/MS data was done using Perseus v2.0.6.0 (Tyanova *et al*., 2016). First, the data was log_2_ transformed obtaining a normal distribution of the data matrix. Then, two-sample Student’s T-tests for significance and fold changes were calculated by comparing each group to the control (i.e. the Milli-Q water at the corresponding timepoint). Principal component analysis (PCA) plots were obtained using Perseus v2.0.6.0 (Tyanova *et al*., 2016). Molecules containing only C, H and O in their formula were retained for these analyses, as they were the ones expected to be generated from PE oxidation.

### Bacterial growth experiments on W-PE

The PE-degrading isolate *Alcanivorax* sp. 24 (Zadjelovic *et al.,* 2020; Zadjelovic *et al.,* 2022), the model alkane degrader *Alcanivorax borkumensis* SK2 (Yakimov *et al.,* 1998), and 14 additional marine isolates from the alkane-degrading taxonomic families *Alcanivoracaceae*, *Marinobacteraceae* and *Halomonadaceae* (Supplementary Table S1) were tested for their ability to grow on W-PE. Overnight cultures were prepared in Marine Broth (MB; BD Difco^TM^). Cells were harvested by centrifuging 1 mL of culture at 16,200 × g for 1 min and washed twice with mineral salts Bushnell–Haas (BH) medium. The BH medium contained (final concentrations): 0.811 mM MgSO₄·7H₂O, 0.136 mM CaCl₂·2H₂O, 7.35 mM KH₂PO₄, 5.74 mM K₂HPO₄, 12.5 mM NH₄NO₃, 0.185 mM FeCl₃·6H₂O, 513.3 mM NaCl, and 0.001% (w/v) yeast extract (Panreac AppliChem), the latter providing essential growth factors for some marine bacteria. The pH was adjusted to 7.2, and the medium was sterilised by autoclaving. Washed cells were inoculated into glass tubes containing 3 mL of BH medium at a starting OD₆₀₀ of 0.01. Disinfected weathered LD-PE fragments at 1 % (w/v, unless stated otherwise) or pyruvate at 0.1 % (w/v) were added as growth substrates. Negative controls included (i) BH medium without added substrate and (ii) uninoculated BH medium containing each substrate to verify sterility. Each condition was tested in triplicate. Cultures were incubated at 30 °C in the dark with orbital shaking at 170 rpm. Growth was monitored by OD₆₀₀ (Ultrospec 1100pro, Amersham Biosciences) at 1, 3, 7, and 14 days. The maximum OD₆₀₀ (maxOD) was defined as the highest value observed across the growth curve. At the end of the experiment, cultures were centrifuged for 5 min at 16,200 ×g. Supernatants (1 mL) were collected, and DOC was measured as described above. The percentage of DOC consumed by each bacterial isolate was calculated using the formula:

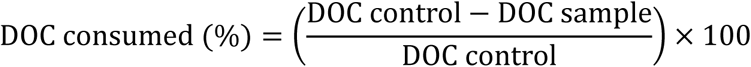

Where ‘DOC control’ was the concentration of DOC measured in the non-inoculated control, representing the initial DOC available and ‘DOC sample’ was the concentration of DOC measured in the inoculated culture at the end of the incubation period.

### Genome sequencing and analysis

Cell biomass from each bacterial isolate (Supplementary Table S1) was harvested from fresh cultures on solid MB medium. Genomic DNA was extracted using the Wizard® Genomic DNA purification kit (Promega) according to the manufacturer’s instructions, and the purified DNA was sent for whole-genome sequencing to MicrobesNG (https://microbesng.com). Libraries were prepared using the Nextera XT Library Prep Kit (Illumina, San Diego, USA), and DNA quantification and normalisation were performed on a Hamilton Microlab STAR automated liquid handling system (Hamilton Bonaduz AG, Switzerland). Sequencing was carried out on an Illumina NovaSeq 6000 platform using a 250 bp paired-end protocol. Adapter trimming of raw reads was performed with Trimmomatic v0.30 (Bolger *et al*., 2014) using a sliding-window quality cutoff of Q15, and *de novo* assemblies were generated with SPAdes v3.7 (Bankevich *et al*., 2012). Assemblies were manually curated to remove spurious contigs, and genome annotation was performed with PROKKA v1.11 (Seemann, 2014). Coding domain sequences (CDS) from all strains used in this study are available as Supplementary Table S2. For taxonomic assignment, whole-genome sequences were submitted to the Type Strain Genome Server (TYGS) for pairwise digital DNA– DNA hybridisation (dDDH) analyses (Meier-Kolthoff and Göker, 2019). Average Nucleotide Identity (ANIb) was calculated with JSpeciesWS (Richter *et al*., 2016) using type strains as references, and these results were visualised as a UPGMA dendrogram constructed in PAST v4.17 (Hammer *et al*., 2001).

To investigate genomic features associated with W-PE assimilation, predicted CDS from each isolate were first combined into a single dataset. The merged sequences were clustered using CD-HIT at 70% sequence identity (Li *et al*., 2001), with sequences within the same cluster considered functionally equivalent. Clusters containing sequences annotated with any of the following keywords were identified: *alkane 1-monooxygenase*, *alcohol dehydrogenase*, *aldehyde dehydrogenase*, *acyl-CoA ligase*, *acyl-CoA dehydrogenase*, *enoyl-CoA hydratase*, *3-hydroxyacyl-CoA dehydrogenase*, and *acetyl-CoA transferase*. The number of sequences within these clusters was used to estimate the gene content associated with each step of the alkane degradation and β-oxidation pathways. Finally, the correlation between gene counts and DOC consumption was assessed by calculating Pearson’s correlation coefficients using PAST v4.17 (Hammer *et al*., 2001).

### High-throughput proteomics

Pyruvate, W-PE and hexadecane grown *Alcanivorax* sp. 24 and *A. borkumensis* SK2 cultures were prepared in triplicate in 3 mL of BH and harvested during the exponential growth phase (OD_600_ of near 0.3). Cells were recovered by centrifugation (16,200 ×g, 1 min at 4 °C) and stored at −20 °C until further analysis. Cell pellets were resuspended in 100 μL of 1×LDS Laemmli buffer (Invitrogen™) containing 1 % β-mercaptoethanol before applying three cycles of thorough vortexing and 5 min incubations at 95 °C. Then, 30 μL of each sample was run for a short migration on a NuPAGE™ 10 % Bis-Tris precast polyacrylamide gel (1.0 mm, 10-well, Invitrogen™) as previously done (Chhun *et al*., 2021). Gel bands containing the entire proteome were cut for further in-gel protein digestion. Gel bands were destained (50 % ethanol and 50 mM ammonium bicarbonate), and standard in-gel reduction-alkylation was performed using 10 mM tris (2-carboxyethyl) phosphine hydrochloride and 40 mM 2-chloroacetamide, after which proteins were in-gel digested overnight with 2.5 ng μL^−1^ of trypsin. The resulting peptide mixture was extracted by sonicating the gel slices in a solution of 5 % formic acid in 25 % acetonitrile, purified with Pierce™ C18 100 μL tips (Thermo Scientific) and finally concentrated at 40 °C in a SpeedVac <u>(Eppendorf® Concentrator Plus)</u> and resuspended in 0.1 % formic acid.

Mass spectrometry analysis was performed using an HRMS timsTOF flex mass spectrometer (Bruker). The resulting peptides were separated on a nanoElute 2 Bruker system. LC-MS/MS analysis was performed by injecting 2 μL onto a precolumn (ThermoScientific PePMap NeO C18, 5μm, 300 μm × 5 mm, 1500 bar) coupled directly to a self-packed analytical column (Ionoptics Aurora Ultimate C18, 1.7 μm, 75 μm × 250 mm, 120 Å, 1000 bar). The chromatography gradient started with an increase of 2 to 25 % mobile phase B (0.1 % formic acid in acetonitrile) in mobile phase A (0.1 % formic acid in water) over 30 min, followed by an increase to 37 % B in 6 min, then 95% B in 1 min, and finally maintained to 95% B during 10 min. Followed by gradient elution into the mass spectrometer (Bruker timsTOF flex MALDI2) equipped with a Bruker Captive Spray source (NSI). A full scan acquisition was performed with a resolution of 35000-40000 mode over a range of 100 to 1700 m/z. The spectrometer was operated in data independent mode (dia-PASEF). The TIMS accumulation time was fixed to 100 ms, and the TIMS separation duration was set to 100 ms. The mobility range was 0.6–1.4 Vs/cm^2^ (1/K0), and the covered m/z range was 200–1000 m/z. The collision energy was distributed from 20 eV to 59 eV.

Raw mass spectral files were processed using Spectronaut® 20 (Biognosys). Spectra were searched using the engine Pulsar. Peptide spectral matches (PSM) to the genome databases were used to calculate a global false discovery rate (FDR). Data were further processed to remove PSMs with an FDR greater than 1.0%. DIA-based protein quantities generated by Spectronaut were further analysed using Perseus v2.0.6.0 (Tyanova *et al*., 2016), with protein signals normalised (i.e. transformed to log_2_, normalised to protein size, and then to total sample signal) prior to comparative proteomic analysis. Polypeptides were considered valid when detected in all three replicates of one condition. Cut-off parameters for comparative analyses used an FDR of 0.01 and S0 (minimal fold change) set to 2. Across pyruvate, hexadecane, and W-PE conditions, 3,378 proteins were detected in *Alcanivorax* sp. 24 and 2,388 in *A. borkumensis* SK2, covering 79.7% and 86.7% of their coding sequences, respectively.

### Metagenomic analysis of polyethylene biofilm microbiomes

Key metabolic pathways implicated in W-PE assimilation were searched for within metagenomes derived from river water microbiomes and from biofilms formed on PE, W-PE, and wood surfaces incubated in the River Sowe, United Kingdom (accession number PRJEB52400, EMBL-EBI; Zadjelovic *et al.,* 2023). Hidden Markov Model (HMMs) profiles used to mine the metagenomes were constructed from the sequences recovered from the UniProtKB database using the keywords: *alkB*; *alkJ*; *adh*; *alkH*; *fadD*; *fadE*, *ACADL*, *ACADM*, *dmdC*; *fadB*, *echA*; and *fadA*. The filter to only obtain bacterial sequences was applied. Up to 200 sequences *per* gene were downloaded from EMBL-EBI and manually curated to exclude incomplete sequences as well as those that did not correspond to the target function.

Metagenome read recruitments were performed with the HMMs for each target gene and for four bacterial house-keeping single copy genes (SCGs; Delmont *et al*., 2018; Sunagawa *et al*., 2013): *ychF* (COG0012), *pheS* (COG0016), *serS* (COG0172) and *leuS* (COG0495). SCGs for HMM construction was obtained from representative NCBI bacterial genomes present in RefSeq V220 (O’Leary *et al*., 2016). Briefly, the first appearing representative genome in the bacteria assembly summary file was chosen for each genus (*n*=3,617) and CDS FASTA files were downloaded. The SCGs from each genome were compiled. Multiple sequence alignments of the W-PE assimilation genes and SCGs were computed using Clustal Omega v1.2.4 (Sievers and Higgins, 2018), and HMMs were subsequently built with hmmbuild within HMMER v3.4 (Eddy, 2011).

HMM validation was performed using hmmsearch on the representative genomes (*n*=3,617) and the genomes of the 17 isolates used in this study (Supplementary Table S1). An e-value cut-off of 10^-25^ was used for SCGs, confirming an expected median of 1 copy *per* genome. The strong correlation between PROKKA-based annotations and HMM-based annotations for the 17 isolates (Supplementary Table S5) confirmed the accuracy of the constructed HMMs.

Metagenomes were mined using hmmsearch for both the W-PE assimilation genes and the SCGs within (i) the raw reads and (ii) the assembled MAGs (completeness >50 % for all). SCG hits (*ychF*, *pheS*, *serS*, *leuS*) with e-value < 10⁻²⁵ were counted *per* sample to establish the baseline estimates of genome (cell) counts. Hits to W-PE assimilation genes with e-value < 10⁻²⁵ were aligned to the reference HMM profiles using hmmalign, and sequences with aligned lengths < 80% of their total length were discarded. We aimed to normalise the results in order to estimate the number of gene copies *per* genome in the community. Results of the reads recruitment were normalised by the alignment length and by the average abundance of the SCGs. For the MAGs, the average abundance of each gene across conditions was calculated by multiplying the number of copies of each gene by the MAG’s relative abundance. Results were then normalised to the average abundance of the SCGs, calculated identically.

## RESULTS

### PE weathering and leachate metabolomic characterisation

Thermal weathering of LD-PE pellets was evident from the characteristic yellowing of the material and was further confirmed by FTIR analysis (Figure 1A). The FTIR spectrum of W-PE exhibited a distinct absorption peak at 1712 cm^-1^, corresponding to the carbonyl (C=O) stretching vibration, a hallmark of polymer oxidation (Gardette *et al*., 2013), which was absent in the pristine PE spectrum. The calculated carbonyl index (CI) of W-PE was 11.43, compared to 0.05 in pristine PE. Additional peaks corresponding to C=C-H bending and C-O stretching were also observed (Figure 1A).

**Figure 1.**
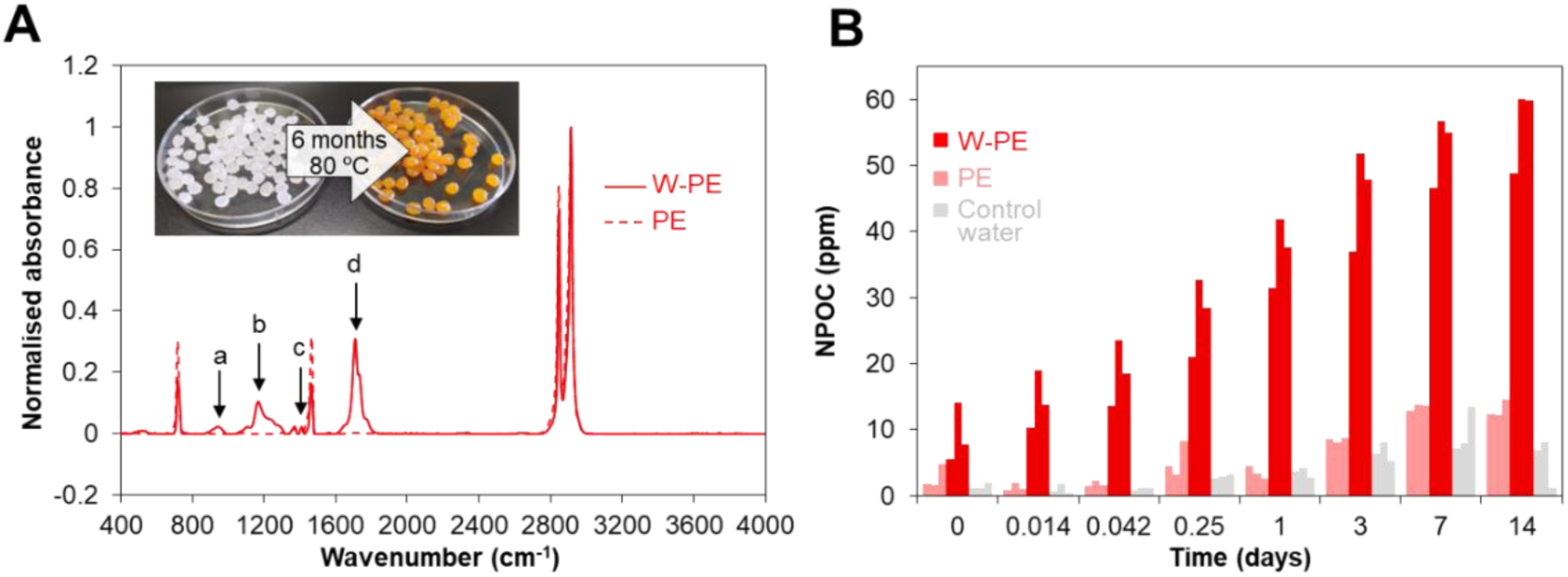
LD-PE pellet weathering and leached DOC. (**A**) FTIR spectra from weathered (thermo-oxidised at 80°C for 6 months; W-PE) and pristine LD-PE pellets (PE). Peaks marked with arrows represent: (a) C=C– H bending, 973 cm^-1^; (b) C–O stretching, 1170 cm^-1^; (c) unaffected region, 1450 cm^-1^; and (d) C=O stretching, 1712 cm^-1^. (**B**) DOC released from 0.3% (w/v) weathered (W-PE) and pristine (PE) LD-PE pellets incubated in sterile Milli-Q water over 14 days, measured as Non-Purgeable Organic Carbon (NPOC). Values of three independent replicates are shown.

As expected, W-PE released substantially more organic carbon into the aqueous phase than pristine PE, with the majority of DOC release occurring within the first 24 h (i.e. accounting for over 60% of the maximum DOC measured by the end the experiment at day 14; Figure 1B). This is consistent with previous reports showing that an initial leaching pulse was followed by slow stabilisation over time (Lee *et al*., 2020; Egea *et al*., 2023). After 14 days, W-PE generated almost 14-fold more DOC than pristine PE, releasing 2.3 ± 0.3% of the polymer’s initial carbon mass as previously reported in a similar experimental setup (Zadjelovic *et al*., 2022). DOC leaching from W-PE was accompanied by strong acidification of the milieu. After 24 h, the pH of the water controls and pristine PE remained at 6.8 ± 0.1 and 7.2 ± 0.1, respectively. In contrast, W-PE caused a pronounced drop in pH to 3.81 ± 0.08, likely due to the accumulation of organic acids generated during polymer oxidation. These pH effects were amplified by using unbuffered Milli-Q water during these experiments.

The metabolomic analysis of the leachate confirmed that random polymer oxidation and polymer scission generated a complex mixture of molecules varying in size and oxidation state. UHPLC-MS/MS identified 2,876 distinct compounds within the DOC fraction released from the plastic (Supplementary Table S3). As expected, while pristine PE released only small amounts of metabolites, clustering closely with water background controls in the PCA, W-PE showed a distinct metabolomic profile from the very beginning of the experiment (i.e. timepoint 0, seconds after water immersion) which became even more evident after 1 h and 7 days (Figure 2A). These results highlight the rapid and extensive release of compounds from W-PE.

**Figure 2.**
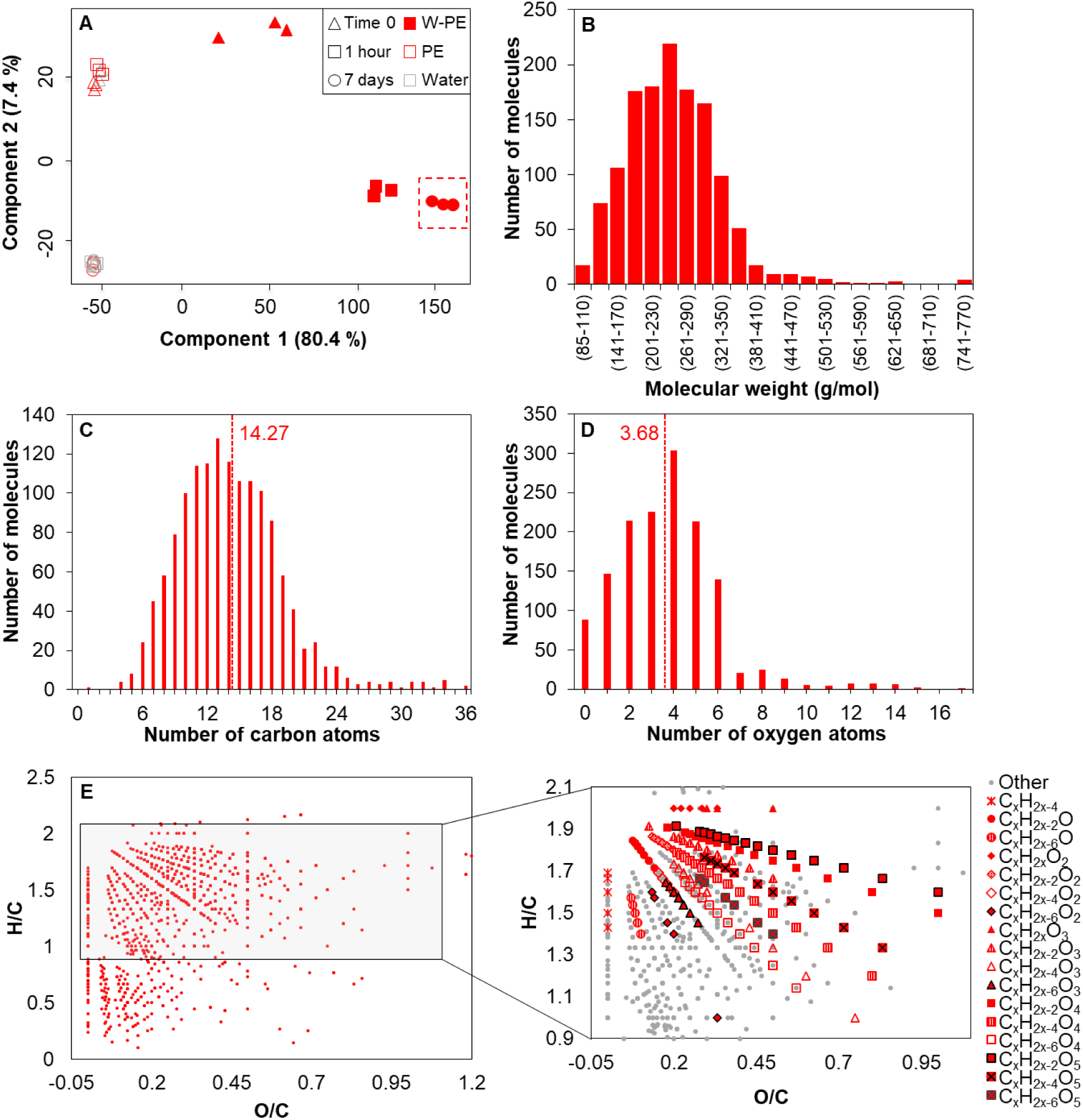
Metabolomic characterisation of W-PE leachate. (**A**) PCA plot based on the UHPLC-MS/MS-detected metabolites within pristine (empty red markers) and weathered PE bead leachates (full red markers), compared to water blanks (empty grey markers), over time. All metabolic data is the result of three independent replicates *per* time and condition. (**B**) Molecular weight distribution of molecules leached by W-PE at timepoint 7 days. Number of carbon (**C**) and oxygen atoms (**D**) in molecules leached by W-PE at timepoint 7 days. Dashed lines in panels C and D indicate the weighted average. (**E**) van Krevelen diagram representing the H/C versus O/C ratios within the detected molecules leached by W-PE at timepoint 7 days. The right panel highlights molecular series with a constant oxidation, but polymer scissions increase by -CH2-units. The plot shows molecules formed only by C, H and O and that were significantly more abundant in weathered plastic samples compared to the background controls after 7 days. Full metabolomic data is available as Supplementary Table S3.

The molecules detected in the W-PE leachate had an average molecular weight of 250.20 ± 81.85 g/mol (Figure 2B). Empirical formula assignments indicated an average chain length of 14 carbon atoms (weighted average: 14.27 ± 5.06; Figure 2C), a size generally considered optimal for bacterial assimilation. Besides possible intermediate oxidation states, the molecules averaged nearly four oxygen atoms *per* molecule (weighted average: 3.68 ± 2.40; Figure 2D). This oxygen content was consistent with PE chain-scission products that had undergone complete oxidation, resulting in dicarboxylic acids. Overall, the average H/C and O/C ratios of the W-PE leachate were 1.37 ± 0.42 and 0.29 ± 0.50, respectively, indicating a high degree of oxidation of these compounds.

To classify the leached molecules, we assumed that oxidation during polymer cleavage occured mainly at chain ends, favouring the formation of carboxylic acids over esters or hydroxy aldehydes. This is consistent with the prevalence of molecules containing four oxygen atoms (Figure 2D), although the assignment remains tentative since no standards were analysed to distinguish isomers. Under this framework, molecular diversity can be simplified as a series differing by –CH₂– units (Figure 2E), as also observed in oxidised HDPE (Eyheraguibel *et al*., 2017), and in agreement with known oxidation mechanisms (Gewert *et al*., 2018). Compounds with H/C > 1.5 likely corresponded to linear saturated alkanols, ketones, aliphatic carboxylic acids, hydroxy-or oxo-carboxylic acids, and dicarboxylic acids with different carbon backbone lengths, while lower H/C ratios indicated unsaturations from extensive oxidation. No formulas matched alkanes (C_x_H_2x+2_) or alkanols (C_x_H_2x+2_O), possibly due to (i) PE polymer scissions do not occur without some kind of terminal oxidation, or (ii) their limited solubility in water. Considering that water-soluble alkanes have been detected in parallel experiments and monounsaturated alkanals (C_x_H_2x-2_O) were predicted in the PE leachates, these observations challenge the latter hypothesis of hydrophobic solubility. Several carboxylic acids were annotated by the prediction software, including 1-nonanoic and decanoic acid, along with three other matching formulas (C_x_H_2x_O_2_). Up to 21 compounds had formulas consistent with monounsaturated carboxylic acids (C_x_H_2x-2_O_2_), including 10-undecenoic acid. However, it should be noted that the same formula could correspond to hydroxy alkanals, such as the annotated 4-hydroxynonenal. In addition, polyunsaturated carboxylic acids (e.g. heptadecatrienoic acid), aliphatic hydroxy acids (e.g. 10-hydroxydecanoic acid), and oxo-carboxylic acids (e.g. 9-oxononanoic acid and 10-oxocapric acid) were identified. The most abundant compound series were those of dicarboxylic acids (C_x_H_2x-2_O_4_), with chain lengths ranging between 5–17 carbons, along with some hydroxy dicarboxylic acids (five oxygen atoms; Figure 2E and Supplementary Table S3).

Overall, the W-PE leachate contained a highly diverse mixture of oxidised aliphatic compounds spanning a wide range of chain lengths, degrees of unsaturation, and functional groups, as shown in Figure 2E. Moreover, these findings challenge the common assumption that PE chain scission primarily generates short-chain alkanes (Wayman and Niemann, 2021); instead, the evidence suggests that terminally oxidised aliphatic compounds are produced, which may subsequently undergo further oxidation to their maximal state, ultimately yielding dicarboxylic acids, as previously observed by Gewert *et al*. (2018). This chemical diversity likely poses a challenge for microbial assimilation, requiring considerable enzymatic versatility to metabolise such structurally heterogeneous substrates.

### Growth and proteomic characterisation of *Alcanivorax* on W-PE

*Alcanivorax* sp. 24, a marine plastic-debris isolate with high PE-degrading potential (Zadjelovic *et al.,* 2020; Zadjelovic *et al.,* 2022), outperformed the model alkane degrader *Alcanivorax borkumensis* SK2 (Schneiker *et al*., 2006; Yakimov *et al*., 1998) in growth and DOC consumption on W-PE leachate (Figure 3). Growth assays in BH mineral medium with varying concentrations of W-PE showed that *Alcanivorax* sp. 24 consistently reached higher OD values and grew faster than SK2 (Figure 3A), although these differences were more pronounced at lower W-PE concentrations (i.e. at 0.15 % W-PE, *Alcanivorax* sp. 24 reached a maximum OD of 0.34 ± 0.01 and SK2 0.13 ± 0.01; 2.53×) than at higher ones (i.e. at 1 % W-PE, *Alcanivorax* sp. 24 grew to 0.62 ± 0.01 and SK2 to 0.40 ± 0.03; 1.56×). Moreover, the growth rate of *Alcanivorax* sp. 24 (1.33 ± 0.14 d⁻¹ in the first 24 h on 1 % W-PE) was nearly double that of SK2 (0.61 ± 0.09 d⁻¹). Such growth differences were specific to W-PE, as no significant difference was observed when both strains were grown on pyruvate (1.77 ± 0.29 d⁻¹ for sp. 24 vs. 1.28 ± 0.36 d⁻¹ for SK2).

**Figure 3.**
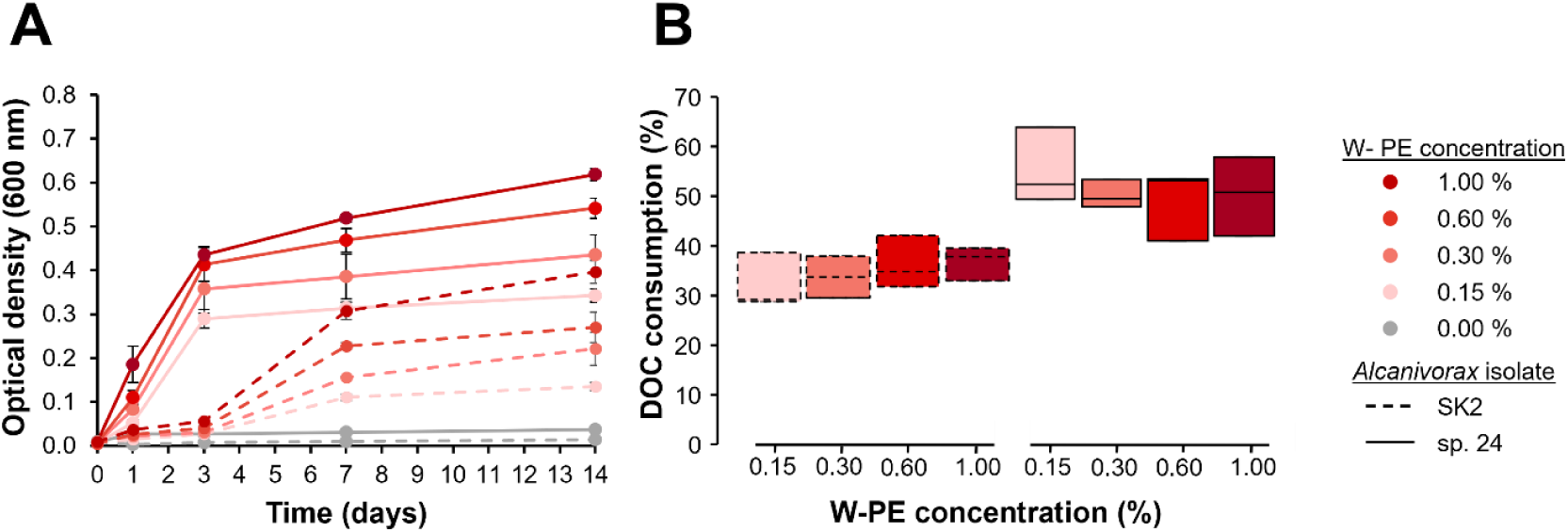
Growth (A) and DOC consumption (B) of *A. borkumensis* SK2 (dashed line) and *Alcanivorax* sp.24 (solid line) on different concentrations of W-PE. Growth curves represent the mean of three biological replicates, with error bars indicating standard deviation. Boxplots of DOC consumption represent values of experimental triplicates, with the median indicated by the black line.

DOC measurements revealed that *Alcanivorax* sp. 24 consumed a significantly higher fraction of organic carbon leached from W-PE than *A. borkumensis* SK2 (50.0 ± 5.8 % *versus* 34.8 ± 4.7 %, *p*-value < 0.001; Figure 3B). Most interestingly, these percentages remained constant regardless of the initial plastic concentration, suggesting that each strain has the metabolic capacity to assimilate a defined fraction of the complex leachate mixture, with *Alcanivorax* sp. 24 displaying a broader assimilation range than *A. borkumensis* SK2. In contrast, both strains consumed similar proportions of the labile substrate pyruvate (87.8 ± 1.4 % DOC consumption for *Alcanivorax* sp. 24 and 85.4 ± 2.0 % for *A. borkumensis* SK2).

The proteogenomic analysis of the two strains revealed that the metabolic differences stem from the higher enzyme redundancy in *Alcanivorax* sp. 24, enabling it to process and assimilate a larger fraction of the complex mixture of oxidised aliphatic compounds present in PE leachates. The established model for aliphatic compound biodegradation involves the stepwise oxidation of molecules with lower oxidation states into fatty acids—typically through sequential reactions catalysed by alcohol and aldehyde dehydrogenases—followed by activation via coenzyme A (CoA) conjugation and subsequent channelling into the β-oxidation pathway (Sabirova *et al*., 2011). While this pathway is relatively straightforward for canonical linear alkanes or fatty acids, the unusual metabolic capacity to assimilate the chemically diverse molecules released from W-PE may arise from either: (i) broad substrate specificity of key enzymes of these pathways, allowing activity on a wide array of compounds, or (ii) the presence of multiple homologous enzymes with overlapping functions but distinct substrate preferences. Comparative proteogenomic analysis between *Alcanivorax* sp. 24 and *A. borkumensis* SK2 supports the latter explanation—a strategy also observed in specialised fatty acid-degrading microbes (Kang *et al*., 2010; Strittmatter et al., 2023; Schiaffi *et al*., 2024). The greater redundancy of key enzymes involved in aliphatic compound metabolism was already apparent at the genomic level. This was particularly pronounced in the early stages of β-oxidation, where *Alcanivorax* sp. 24 encodes markedly higher numbers of acyl-CoA ligases (7.8×), acyl-CoA dehydrogenases (2.1×), and enoyl-CoA hydratases (2.5×) compared to SK2 (Supplementary Table S2).

Shotgun proteomics confirmed that the broader metabolic potential of *Alcanivorax* sp. 24 was not only genomically encoded but also overproduced in the presence of W-PE. This feature was not seen when grown with hexadecane (Figure 4). When considering only proteins significantly overproduced in hexadecane when compared to the pyruvate control condition, both *Alcanivorax* strains produced a reduced number of canonical enzymes to funnel the alkane through the initial oxidation steps, CoA ligation and final β-oxidation cycle (right side of Figure 4). Other than the acyl-CoA ligase and hydratase/dehydrogenase steps, where *Alcanivorax* sp. 24 had approximately twice as many enzymes, the two strains showed comparable enzyme counts across the remaining steps. Conversely, when grown on W-PE, *A. borkumensis* SK2 displayed enzyme numbers largely consistent with those detected on hexadecane, whereas *Alcanivorax* sp. 24 exhibited a marked increase in both the number and overexpression of enzymatic isoforms, particularly for the CoA ligase and β-oxidation steps (left side of Figure 4).

**Figure 4.**
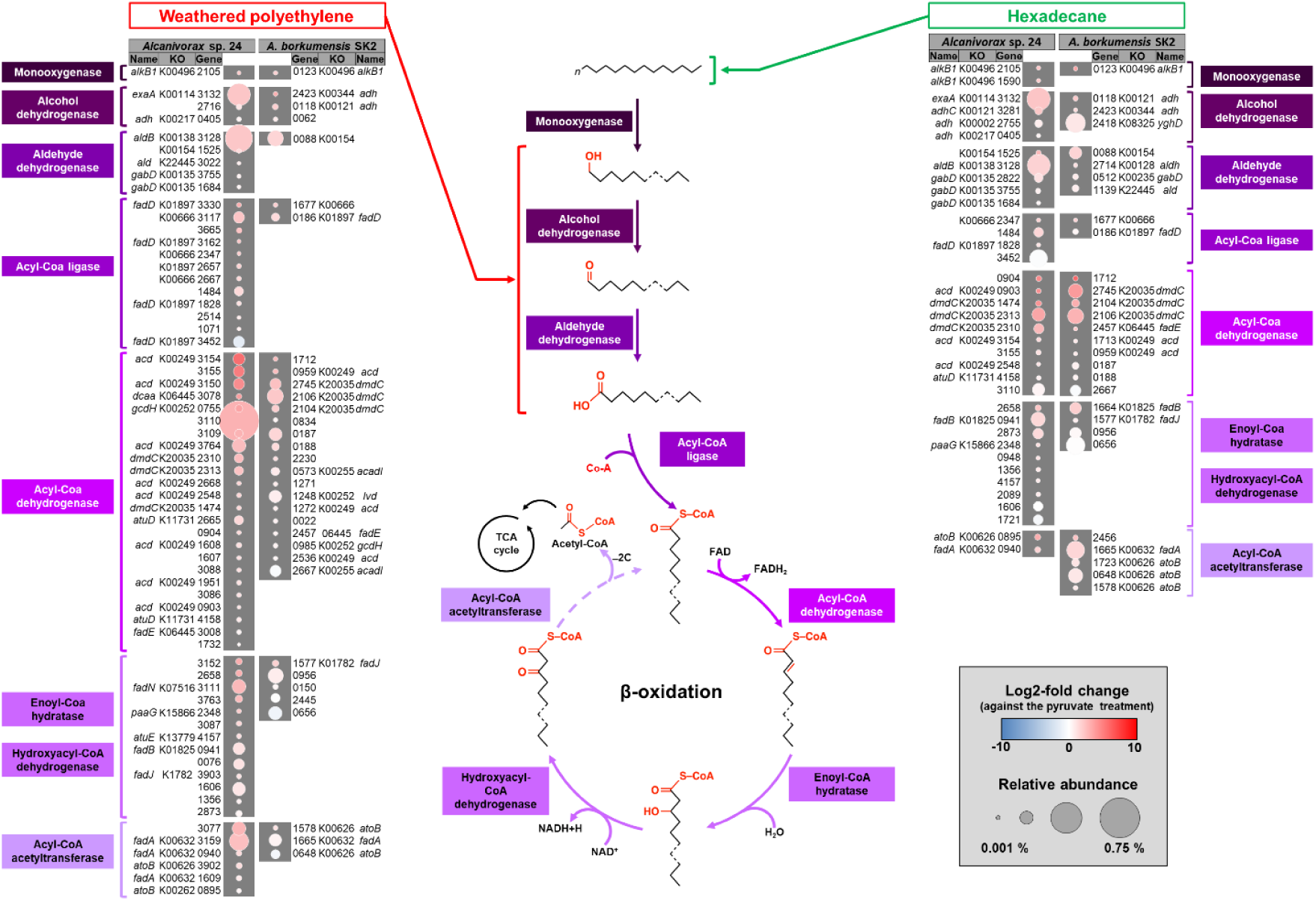
Proteomic response of *Alcanivorax* sp. 24 and *A. borkumensis* SK2 grown with W-PE or hexadecane. Only proteins involved in the metabolism of aliphatic substrates and that were significantly overproduced relative to the pyruvate control are shown. Relative abundance represents the average of three independent biological replicates. Complete proteomic data is available as Supplementary Table S4.

To better understand this process, we next examine each enzymatic step in detail:

1. Alkane monooxygenase. The initial oxidation step for alkanes is catalysed by alkane 1-monooxygenases (AlkB). In *Alcanivorax* sp. 24, the isoenzyme ALC24_2105 was abundant in both, W-PE and hexadecane, conditions, whereas a second AlkB ALC24_1590 was overproduced only with hexadecane (Figure 4). This initial step could also be performed by two cytochrome P450s (ALC24_2795 and ALC24_1244), which were detected but not overproduced. By contrast, the subterminal monooxygenase ALC24_0335, known as the Baeyer-Villiger oxidation, was specifically overproduced only in the presence of W-PE, suggesting a role in secondary hydroxyl metabolism. *A. borkumensis* SK2 encodes a single AlkB (ALCSK2_0123) and two cytochrome P450s (ALCSK2_0203, ALCSK2_2390) –all overproduced in hexadecane and W-PE– but lacks any Baeyer-Villiger monooxygenases. The reduced AlkB production observed in both *Alcanivorax* strains when exposed to W-PE likely reflects the absence of hexadecane or similar medium-chain alkanes, which typically induce AlkB-related monooxygenases.
2. Alcohol dehydrogenase (ADH). Different ADHs classes can be involved in the oxidation of aliphatic alcohols, including NAD(P)^+^-dependent, quinoproteins, zinc-dependent, and iron-containing forms, without a clear pattern of their specificity across substrates. In *Alcanivorax* sp. 24, the quinoprotein ALC24_3132 was consistently the most abundant whatever the condition, while the zinc-dependent ALC24_3281 and NAD(P)^+^-dependent ALC24_2755 were hexadecane-specific, and the zinc-dependent ALC24_2716 was specific to W-PE. In *A. borkumensis* SK2, the zinc-dependent ADH ALCSK2_0118 and NAD(P)^+^-dependent ALCSK2_2423 were expressed at similar levels in both conditions, whereas the NAD(P)^+^-dependent ALCSK2_2418 was more abundant in hexadecane.
3. Aldehyde dehydrogenases (ALDH). Also, different types of ALDHs can participate in the metabolism of aliphatic compounds. While *Alcanivorax* sp. 24 abundantly expressed the NAD(P)^+^-dependent ALC24_3128 in both conditions, the broad-range NAD(P)^+^-dependent ALC24_3022 was specific to W-PE, and the NAD-dependent ALC24_2822 was hexadecane-specific, though both were detected at low levels. In *A. borkumensis* SK2, only the NAD(P)^+^-dependent ALCSK2_0088 was overproduced in W-PE, while three additional ALDHs were enriched in hexadecane.
4. Other oxidoreductases (Supplementary Table S4). Apart from the already highlighted enzymes, several additional oxidoreductases representing over 0.1 % of the proteome of *Alcanivorax* sp. 24 were detected when grown with W-PE, including ALC24_3140 (14× fold-change), ALC24_1446 (6×), ALC24_3765 (5×), ALC24_3766 (5×), and ALC24_3146 (4×). Others were strongly overproduced, such as ALC24_2300 (33×), ALC24_0630 (15×), ALC24_2036 (14×), ALC24_3158 (8×), and ALC24_3148 (7×). On the other hand, when grown with hexadecane, only ALC24_2335 (2.4×) and ALC24_2568 (1.5×) exceeded 0.1 % of the proteome, with a handful of others moderately enriched (Supplementary Table S4). In *A. borkumensis* SK2, three uncharacterised oxidoreductases (ALCSK2_1028, 24.7×; ALCSK2_2045, 5.4×; ALCSK2_0086, 3×) were abundant in W-PE, whereas ALCSK2_2399 (18×) was more enriched in hexadecane.
5. Acyl-CoA ligase (ACAL). The most considerable differences between isolates were clearly observed in isoforms of ACALs. In *Alcanivorax* sp. 24, most ACALs were strongly overproduced in the presence of W-PE, while only three (ALC24_2347, FACS; ALC24_1484, long-chain FACS; ALC24_1828, ACDL/FadD) were expressed at similar levels across conditions. Others, including ALC24_3665 (AMP-binding, medium-chain) and ALC24_3330 (ACDL/FadD), were W-PE-specific. *A. borkumensis* SK2, in contrast, produced only two ACALs (ALCSK2_1677, FACS; ALCSK2_0186, ACDL/FadD), with only a slightly higher abundance in W-PE.
6. Acyl-CoA dehydrogenase (ACAD). Both isolates produced the same number of ACADs under hexadecane (*n*=10). However, W-PE induced the production of six additional ACADs in *Alcanivorax* sp. 24 compared with SK2, and at higher abundances. Many were specific to W-PE and were represented by different KOs, consistent with a broader substrate range. The highly abundant ALC24_3110 appears to be a pimeloyl-CoA dehydrogenase, an enzyme required for the processing of dicarboxylic acids. *A. borkumensis* SK2 did not show such condition-dependent changes. Interestingly, both isolates overproduced three 3-methylmercaptopropionyl-CoA dehydrogenases (DmdC), which is usually associated with sulphur metabolism but may also be involved in fatty acids metabolism.
7. Final steps of the β-oxidation pathway. *Alcanivorax* sp. 24 also produced more enoyl-CoA hydratases than *A. borkumensis* SK2 in W-PE, with higher relative abundance. By contrast, the canonical FadB homologs ALC24_0941 in *Alcanivorax* sp. 24 and ALCSK2_1664 in *A. borkumensis* SK2 were more abundant under hexadecane. Finally, in *Alcanivorax* sp. 24, the two most abundant Acyl-CoA acetyltransferases in W-PE were not detected in hexadecane, highlighting a clear substrate-specific adaptation at the terminal step of β-oxidation.

Altogether, the proteomic data indicated metabolic redundancy as a key strategy for chemically funnelling the diverse array of compounds released from W-PE into central metabolism, thereby maximising metabolite assimilation. Although the precise roles of many non-canonical enzymes remain unclear, their strong induction in the presence of W-PE suggests previously unexplored functions in the metabolism of alkanes and polymer-derived compounds.

### Metabolic redundancy positively correlates with PE mineralisation in other bacterial groups

The growth of 17 marine isolates from three taxonomic groups (*Alcanivoracaceae*, *Halomonadaceae*, and *Marinobacteraceae*) was monitored on W-PE for 14 days, after which DOC consumption was quantified. Growth rates, maximum OD, and DOC utilisation varied widely both between and within each group (Figure 5A). The two new *Alcanivoracaceae* strains added to this latest analysis, i.e. *Alloalcanivorax dieselolei* B5 and *Alloalcanivorax xenomutans* JC109, showed slightly lower but comparable growth rates and maximum OD to *Alcanivorax* sp. 24. Within *Halomonadaceae*, we observed both the fastest and slowest growth rates, closely correlated–as expected–with maximum ODs, underscoring the heterogeneity within this group. In contrast, *Marinobacteraceae* isolates displayed both lower growth rates and lower maximum OD values than the other groups (Figure 5A). The most striking observation emerged from DOC consumption. While expected higher growth yields positively correlated with greater DOC utilisation in *Alcanivoracaceae* and *Halomonadaceae*, no such relationship was observed in *Marinobacteraceae* (Figure 5B). Despite the consistent low growth yields across strains, DOC consumption amongst *Marinobacteraceae* ranged widely (−8.24% to 54.81%), indicating that increased substrate uptake did not translate into increased biomass in these isolates. This discrepancy may have two possible explanations: (i) technical limitations of OD-based planktonic growth measurements (e.g. differences in cell size, aggregation or biofilm formation), although this seems unlikely since aggregation or biofilm formation was only seen in *M. nauticus* V40; or (ii) a physiological strategy in which some *Marinobacteraceae* isolates channel PE leachates as a source of energy or carbon storage rather than a carbons source for growth. Growth limitation caused by the medium’s lack of growth factors was unlikely, as all *Marinobacteraceae* isolates grew well on pyruvate, reaching OD > 0.480.

**Figure 5.**
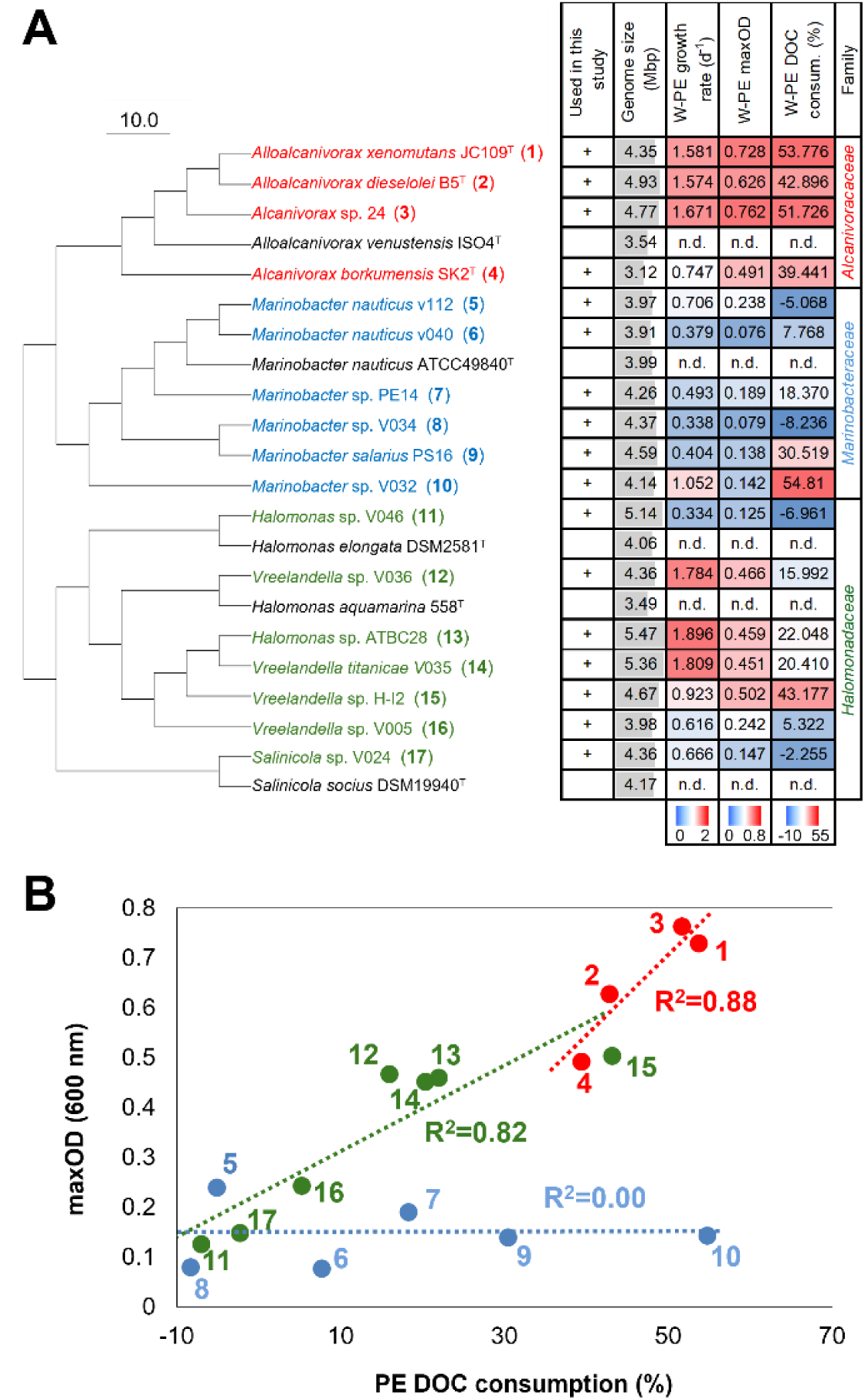
Assessment of W-PE assimilation amongst bacterial isolates from the families *Alcanivoracaceae* (red), *Halomonadaceae* (green) and *Marinobacteraceae* (blue). (**A**) UPGMA dendrogram based on ANIb values of five type strains and the 17 isolates used in this study (indicated by ‘+’). For each isolate, genome size (Mbp), growth rate (d^-1^), maximum optical density (OD600 ‘maxOD’), and percentage of dissolved organic carbon (DOC) consumed after 14 days of growth with 1% W-PE are shown. (**B**) Correlation between the maximum OD600 reached by each strain and the percentage of W-PE DOC consumed after 14 days. Numbers in panel B correspond to the strain identifiers shown in panel A.

Genomic analysis of the 17 isolates revealed that the ability to metabolise W-PE leachates correlated more with the redundancy within the β-oxidation pathway than with alkane-degradation enzymes (Figure 6). Genes from all 17 isolates were clustered at >70% amino acid identity, assuming a shared function within each cluster (Tian and Skolnick, 2003). Annotations were then examined for key steps in alkane oxidation and fatty acid β-oxidation (Supplementary Table S2). To assess the contribution of each step, the number of gene copies *per* enzyme in each isolate was correlated with its DOC consumption from W-PE (Figure 6). A positive correlation would suggest that redundancy at a given step broadens substrate assimilation, thereby increasing PE-derived DOC consumption. Only a weak correlation was detected for *alkB* (R² = 0.33, P = 0.016), with no significant relationships for other alkane-oxidation genes (*alkJ*, *alkH*), either individually or combined (Figure 6). Moreover, *alkB* proved a poor predictor of PE degradation: *Vreelandella* sp. H-I2 consumed 43% of W-PE DOC despite lacking *alkB*, whereas *Marinobacter* sp. V034, carrying two monooxygenases, showed the lowest leachate consumption (−8.24%). By contrast, strong positive correlations emerged between DOC consumption and gene redundancy across β-oxidation steps (Figure 6). This relationship was mainly driven by the three *Alcanivoracaceae* strains, which consumed the highest DOC fractions and carried the greatest number of isoenzymes *per* catabolic step. Amongst these, the first and last step of β-oxidation–i.e. acyl-CoA dehydrogenase and acyl-CoA acetyltransferase, respectively–showed the strongest correlation (R² = 0.49, P = 0.0019 and R² = 0.57, P = 0.0005). A similar relationship was found when considering the entire β-oxidation pathway (R² = 0.48, P = 0.0021), highlighting the key role of β-oxidation redundancy in channelling the chemically diverse aliphatic derivatives generated during PE weathering into central metabolism.

**Figure 6.**
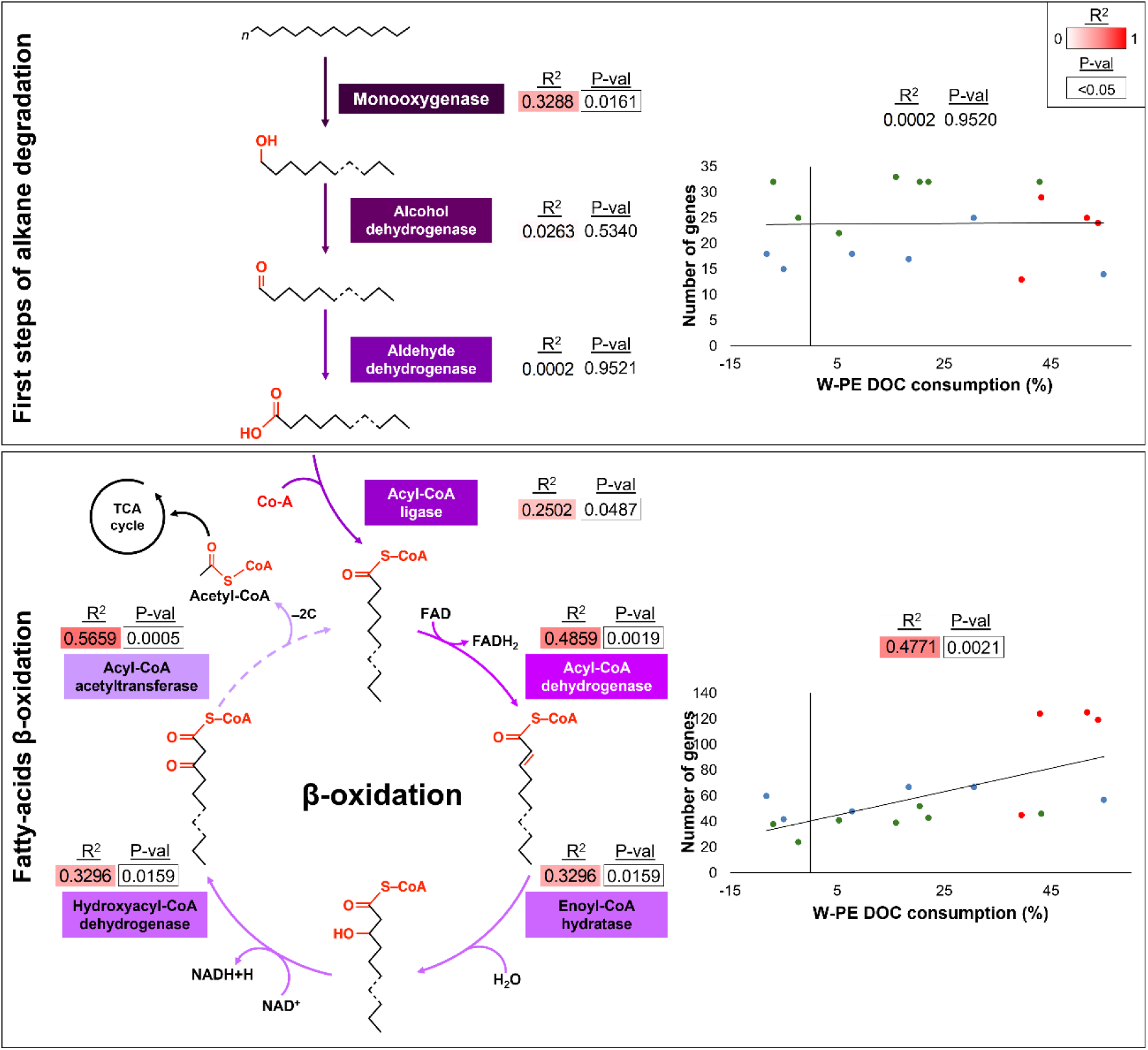
Correlation between W-PE DOC consumption and the gene copy number of each step within the alkane degradation and fatty-acid β-oxidation pathways of 17 PE-degrading isolates. Pearson correlation for each metabolic step and the overall pathway is shown. Isolates from the taxonomic groups *Alcanivoracaceae* (red), *Halomonadaceae* (green) and *Marinobacteraceae* (blue) are represented.

### Enrichment of β-oxidation pathway genes plastisphere metagenomes from PE and W-PE plastic films

The metagenomic analysis of microbiomes from pristine PE and W-PE films incubated for seven days in the River Sowe (UK) revealed a strong enrichment of alkane- and fatty-acid–degradation genes compared with the surrounding river water, and a weaker—but still statistically significant—enrichment relative to biofilms formed on wood (Figure 7). To contextualise these findings, we first compared the number of gene copies in a set of randomly selected representative genomes (*n* = 3,617) with those in the genomes of the 17 PE-degrading isolates used in this study. As expected, the PE-degrading isolates contained, on average, 3-fold more copies of metabolic genes than the randomly selected genomes (Figure 7A).

**Figure 7.**
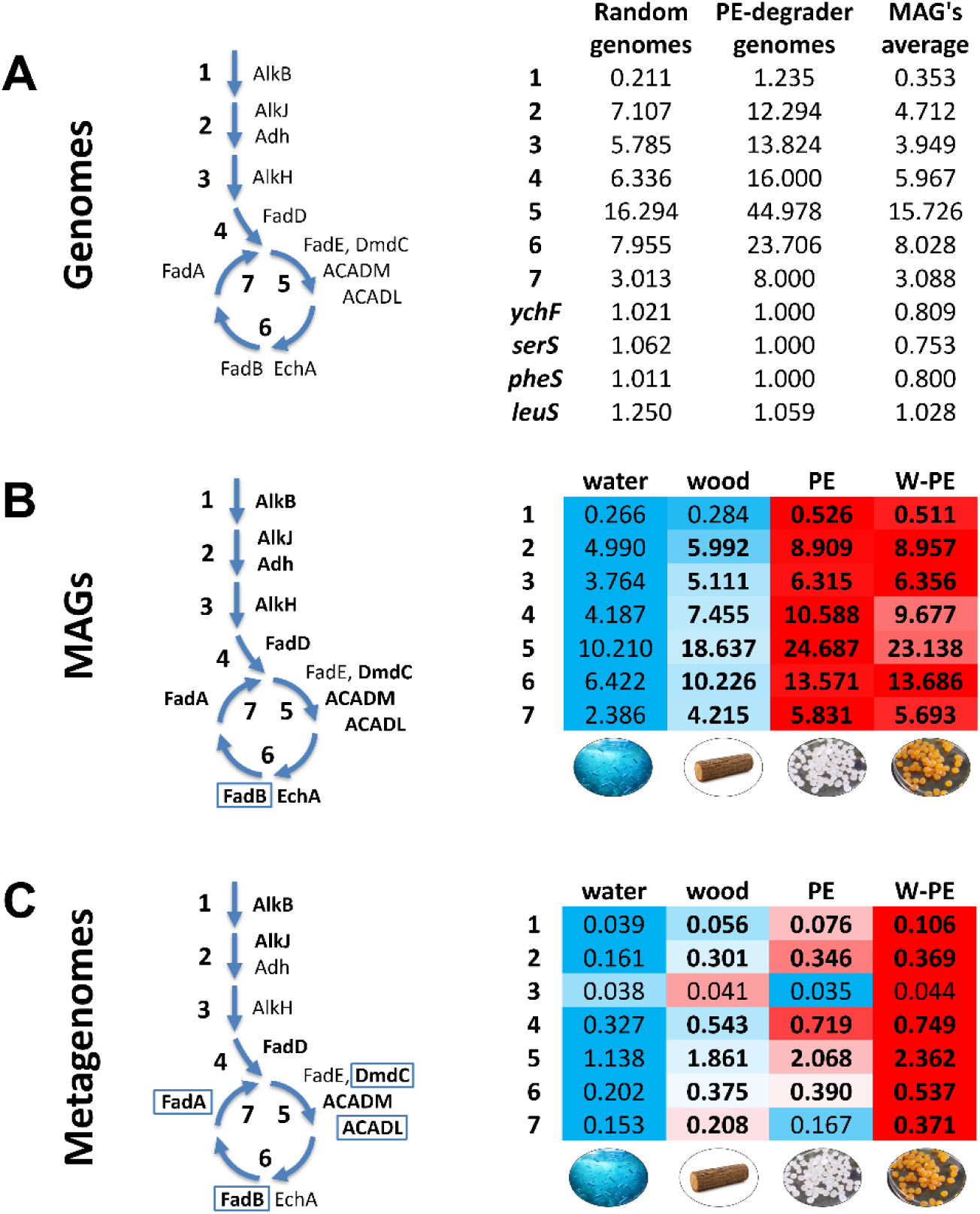
Redundancy of alkane degradation and fatty-acid β-oxidation pathways in metagenomes of PE plastispheres. Metagenomes were obtained from pristine and weathered low-density PE films (*PE* and *W-PE*), and from wooden strips used as a control surface (*wood*). These substrates were incubated *in-situ* for seven days in the River Sowe (Coventry, UK; Zadjelovic *et al*., 2023) after which total DNA was extracted and sequenced, alongside the surrounding planktonic microbial community (*water*). Control genomes, MAGs and metagenomes were searched using HMM profiles for each gene involved in the seven steps of the metabolic pathways of interest, as well as for single-copy housekeeping genes (*ychF*, *serS*, *pheS* and *leuS*). (**A**) Average gene copy number for each metabolic step and housekeeping genes across 3,617 representative genomes from NCBI (*random genomes*), 17 genomes from bacteria that consumed large amounts of organic carbon leached from W-PE (*PE-degrader genomes*), and 215 MAGs generated from the metagenomic dataset. (**B**) Average gene copy number *per* MAG, weighted by their relative abundance in each sample and normalised using the single-copy housekeeping genes. (**C**) Average gene copy number *per* cell in each metagenome using a read-based approach, corrected for gene length and normalised using the single-copy housekeeping genes. Gene copy numbers shown in bold indicate significant increases relative to the water control (p < 0.05). The heatmap displays the lowest (blue) and highest (red) gene copy numbers withing each step, with the median (percentile 50) shown in white. In the schematic pathway, genes shown in bold were significantly enriched in W-PE biofilms compared with both water and wood controls (p < 0.05). In contrast, genes marked with a square were also significantly more abundant in W-PE than in PE (p < 0.05). Values represent the average of three biological replicates.

Interestingly, the assembled MAGs obtained from the planktonic and biofilm microbiomes displayed average gene copy numbers similar to those observed across the randomly selected representative genomes (Figure 7A), suggesting that PE-degrading bacteria are sparsely represented amongst the 215 MAGs. Nevertheless, when considering the abundance of these MAGs across conditions, the HMM-based analyses showed that nearly all analysed genes increased significantly in abundance in PE and W-PE plastispheres relative to the surrounding water (2.1×) and wood (1.4×) controls (Figure 7B). The only exception was *fadE*, which was less abundant and generally rare across MAGs (Supplementary Table S5). Although the abundance of redundant genes within each metabolic step in PE and W-PE biofilm microbiomes exceeded that observed in the representative genomes, it did not reach the elevated values characteristic of the PE-degrading isolate genomes (Figure 7A and 7B). This pattern suggests that PE-degrading bacteria—encoding high numbers of redundant alkane- and fatty-acid–degradation genes—likely constitute only a minority of the plastisphere community, and that their signal is diluted by the presence of accompanying non-PE-degrading microbes within these biofilms.

The genome-centric analysis of the metagenomes also revealed clear taxonomic shifts amongst biofilms formed on PE, W-PE, and wood, as well as between these biofilms and the planktonic microbiome from the surrounding river water. All biofilms showed strong enrichment of members of the *Burkholderiaceae* family (71.35 %, 66.09 % and 49.43 % in PE, W-PE and wood, respectively), followed by *Pseudomonadaceae* (15.55 %) on W-PE, *Moraxellaceae* on pristine PE (9.86 %), and *Flavobacteriaceae* and *Rhodocyclaceae* on wood (7.11 % and 4.37 %, respectively). Families with the highest average numbers of alkane- and fatty-acid–degradation genes included *Saprospiraceae* (162 average sum of degradation genes), *Polyangiaceae* (90), *Microbacteriaceae* (75), *Hyphomonadaceae* (69), *Pseudomonadaceae* (68), *Sphingomonadaceae* (67), *Rhodobacteraceae* (67), *Nannocystaceae* (65), *Burkholderiaceae* (64), and *Turneriellaceae* (61). Intriguingly, MAGs from *Saprospiraceae* (2 MAGs) and *Polyangiaceae* (4 MAGs) were more enriched in wood biofilms, despite their high gene copy numbers. *Rhodobacteraceae* (2 MAGs) and *Pseudomonadaceae* (3 MAGs) were the only of these families whose MAGs encoded, on average, two copies of *alkB*. The most abundant MAG in PE and W-PE plastispheres, *Sphaerotilus natans* (MAG 180; 32 % in W-PE and 22 % in PE), did not exhibit elevated numbers of alkane- or fatty-acid–degradation genes and, although it encoded the main steps of β-oxidation, it lacked *alkB*, suggesting that it may metabolise pre-oxidised intermediates rather than initiate alkane activation. By contrast, MAG 168 (unclassified) carried the highest total number of targeted genes across all MAGs yet did not show high abundance on any substrate. A similar pattern was observed for *Haliscomenobacter hydrossis* (MAG 153), a representative of *Saprospiraceae* and a known organic-matter degrader: despite harbouring many relevant genes, it was found predominantly on wood. Conversely, two *Burkholderiaceae* genomes (MAG 74 and MAG 99) combined high fatty-acid–degradation gene copy numbers with moderate enrichment in PE and W-PE biofilms (1.22 % and 0.42 %, respectively, against 0.03 % in water for MAG 74 and 1.83 % and 0.70 % against 0.05 % in water for MAG 99), driven in particular by elevated numbers of *fadD* genes.

We also performed a raw read–based analysis of the metagenomes (Figure 7C), which overall yielded lower metabolic gene copy numbers than those obtained from the isolate genomes and MAGs. This likely reflects the technical limitations of searching short-read metagenomes with HMMs: whereas highly conserved housekeeping SCGs recruit reads efficiently, many metabolic genes are more divergent, resulting in poorer recruitment and consequently lower estimated gene copy numbers after SCG normalisation. Despite this overall reduction, the fold-change increase observed in W-PE relative to the other control conditions closely matched the pattern recovered in the genome-centric MAGs analysis. Specifically, W-PE plastispheres showed a significant enrichment of alkane- and fatty-acid–degradation genes—2.2× higher than in water and 1.4× higher than in wood (Figure 7C). Interestingly, this analysis also revealed a higher abundance of these genes in W-PE compared with pristine PE plastispheres (1.4×), consistent with the expectation that readily available weathered substrates promote faster growth of PE-assimilating bacteria—an effect less pronounced on the more recalcitrant pristine polymer.

## DISCUSSION

This study highlights the central role of the fatty acid degradation pathway in the metabolisation of W-PE. Using a multi-layered approach—including the detailed proteomic characterisation of two *Alcanivorax* isolates, growth assays of 17 isolates linked to their genomic features, and metagenomic analyses of PE and W-PE colonising communities—we provide new insights into the molecular basis of plastic-associated carbon assimilation. Previous studies reported that neither *Alcanivorax* sp. 24 nor *A. borkumensis* SK2 showed substantial growth on pristine LD-PE as a sole carbon and energy source, although slow biodegradation was suggested (Rose *et al*., 2020; Zadjelovic *et al*., 2022). For this reason, we focused on W-PE, exploring the metabolic pathways required for the assimilation of its leachates rather than the bio-oxidation of the intact polymer, which is a fundamentally different process.

Some parallels can be drawn between alkane and W-PE assimilation, though clear distinctions exist. For example, *A. borkumensis* SK2 growing on hexadecane overexpressed only one acyl-CoA dehydrogenase (ABO_1653) and one enoyl-CoA hydratase (ABO_2102; Sabirova *et al*., 2011), with similar observations by Naether *et al*. (2013). In contrast, *Alcanivorax* sp. 24 showed overproduction of a much broader set of enzymes, including up to 13 enoyl-CoA hydratases, 11 acyl-CoA dehydrogenases, and 3 acetyl-CoA transferases in response to alkanes and medium-chain dicarboxylic acids (Zadjelovic *et al*., 2022). This supports the idea that SK2 is specialised for linear alkane degradation, while isolate 24 is more versatile and capable of funnelling a wider array of molecules—such as those leached by W-PE—as confirmed by the present results. Our proteogenomic analyses further demonstrated the overproduction of enzymes linked to fatty acid degradation, enabling growth on W-PE.

Enzyme redundancy within the β-oxidation pathway appears crucial for a more efficient W-PE assimilation. However, precisely assigning functions to individual enzymes was not feasible, highlighting the need for further physiological and enzymatic studies. Identifying the specific leachate molecules assimilated by each strain and linking them to enzyme specificities would be particularly valuable. The high degree of redundancy complicates functional characterisation through knockout mutagenesis (Schiaffi *et al*., 2024), limiting our ability to resolve the precise contribution of each enzyme. In addition, other genomic features may contribute to the process, such as auxiliary enzymes (e.g. isomerases, oxidoreductases) and diverse transporters with poorly defined substrate specificities (van der Hoek and Borodina, 2020).

Our findings also challenge the assumption that the presence of hydrocarbonoclastic bacteria directly indicates PE degradation capacity. Although hydrocarbon degraders are frequently detected in plastisphere communities (Delacuvellerie *et al*., 2019; Pinto *et al*., 2020), the differential abilities of closely related taxa, such as *Alcanivorax* sp. 24 and *A. borkumensis* SK2, demonstrate that taxonomic affiliation alone is not predictive of degradation potential. Therefore, community-level claims of plastic biodegradation should be supported by genetic and functional evidence rather than taxonomy alone. Gene abundance must also be considered, as the redundancy of catabolic genes strongly influences metabolic potential, as we demonstrate in this study.

Previous studies with microbial communities reported rapid consumption of DOC leached from plastics: up to 85 % within 16 days (Egea *et al*., 2023) and about 60 % within 5 days (Romera-Castillo *et al*., 2018). These communities often resembled those responding to natural marine DOM, dominated by taxa such as *Alteromonas*, other Gammaproteobacteria, and Roseobacter (Birnstiel *et al*., 2022). However, these experiments used commercial plastics, where leachates were likely dominated by additives rather than weathered polymer byproducts. In our study, the maximum DOC consumption by a single isolate was 54.8 %, indicating that many leachate molecules remain inaccessible to individual strains. This suggests that microbial consortia with complementary metabolic capabilities may be necessary for more complete W-PE utilisation.

Metagenomic analyses of PE- and W-PE plastispheres revealed a significant enrichment of both alkane- and fatty-acid–degradation pathways compared with the microbiomes of wood biofilms and surrounding river water. Although both complementary HMM-based search approaches (raw read recruitment and genome-centric analysis using MAGs) detected significant enrichment, the overall redundancy of these pathways in PE plastispheres remained far below that expected for PE-degrading isolates and, instead, resembled the gene copy numbers found in average reference genomes. This pattern reflects the essentially stochastic colonisation of plastics and the low representation of true degraders—a phenomenon that may be exacerbated in mature biofilms, where biodegraders can represent an even smaller relative proportion of the community (Erni-Cassola *et al*., 2020; Purganan *et al*., 2025). The concurrent enrichment of alkane- and fatty acid-degradation pathways should be interpreted with caution. Microbes possessing highly redundant β-oxidation pathways—optimal for assimilating W-PE leachates composed of diverse oxidised aliphatic molecules—are often, but not always, also equipped with robust alkane-degradation pathways. Here we show that some efficient W-PE leachate consumers lack alkane monooxygenase (*alkB*), indicating that enrichment of *alkB* does not necessarily reflect PE degradation directly but may arise from its co-occurrence with fatty-acid degradation genes.

The success of several abundant plastisphere members was likewise not explained by high redundancy in β-oxidation genes. Instead, traits such as strong biofilm-forming capacity or the ability to exploit metabolites oxidised by other community members may underlie their competitive advantage. Conversely, some taxa with extensive PE-assimilation potential did not specifically thrive on plastics, indicating that metabolic capacity alone does not determine substrate colonisation; ecological interactions, niche availability, and metabolic cross-feeding are likely to play important roles (Zhao *et al*., 2024). Nevertheless, the significant enrichment of alkane- and fatty-acid–degradation pathways in W-PE plastispheres relative to surrounding water microbiomes (2.2×) and wood biofilms (1.4×) underscores the importance of metabolic redundancy in enabling microbial communities to collectively assimilate a greater fraction of the complex compounds present in PE leachates. Ultimately, confirming the functional relevance of these genes will require evidence of gene expression and enzyme abundance as done here with monocultures. Finally, although *fadE* is considered the canonical β-oxidation dehydrogenase, our data demonstrate that other dehydrogenases—such as long- and medium-chain acyl-CoA dehydrogenases (*ACADL*, *ACADM*) and the broad-specificity linear acyl-CoA dehydrogenase *dmdC*—play a more prominent role during W-PE assimilation.

## CONCLUSIONS

This study demonstrates that bacterial assimilation of W-PE leachates relies primarily on the redundancy and versatility of its fatty-acid degradation pathway, rather than on narrow specialisation alone. While some organisms—such as *Alcanivorax borkumensis* SK2—appear highly specialised for linear alkane metabolism, others, including *Alcanivorax* sp. 24, possess far more versatile and redundant metabolic repertoires that enable the assimilation of the chemically diverse oxidised aliphatic compounds released from W-PE. These findings expose the limitations of inferring plastic-degrading potential from taxonomy alone and emphasise the need to integrate genomic, proteomic, and functional analyses when assessing microbial contributions to plastic mineralisation. The incomplete utilisation of leachates by individual isolates further suggests that microbial consortia with complementary metabolic capacities may be required for more efficient W-PE assimilation, consistent with the inherently complex and heterogeneous nature of PE-derived substrates. Although alkane- and fatty-acid–degradation pathways emerged as key and consistently enriched mechanisms, additional metabolic routes, transport systems, and interspecies interactions likely shape overall community performance and remain underexplored. Collectively, this work deepens our understanding of microbial metabolism of plastic-derived organic carbon and underscores the need for integrative, multi-omic studies to elucidate the mechanisms governing plastic assimilation in natural environments. Notably, the analytical framework established here provides robust tools for future microbiome-based searches for genes, pathways, and taxa involved in PE assimilation, setting the stage for more accurate predictions of plastic turnover potential across ecosystems.

## SUPPLEMENTARY INFORMATION

Additional file 1: Supplementary Table S1. List of marine isolates from the alkane-degrading taxonomic families *Alcanivoracaceae*, *Marinobacteraceae* and *Halomonadaceae*.

Additional file 2: Supplementary Table S2. Coding Domain Sequences (CDS) of the *Alcanivorax* sp. 24*, Alcanivorax borkumensis* SK2, *Alloalcanivorax dieselolei* B5, *Alloalcanivorax xhenomutans* JC109, *Halomonas* sp. ATBC28, *Halomonas* sp. V036, *Halomonas* sp. V046, *Marinobacter* sp. PE14, *Marinobacter* sp. V032, *Marinobacter* sp. V034, *Marinobacter nauticus* V040, *Marinobacter nauticus* V112, *Marinobacter salaries* PS16, *Salinicola* sp. V024, *Vreelandella* sp. H-I2, *Vreelandella* sp. V005, *Vreelandella titanicae* V035, genomes.

Additional file 3: Supplementary Table S3. Metabolomic analysis of PE and W-PE leachates at time points 0, 1h and 7 days after submersion in water.

Additional file 4: Supplementary Table S4. Proteomics analysis and comparative proteomics of *Alcanivorax borkumensis* SK2 and *Alcanivorax* sp. 24 when grown with pyruvate, W-PE and hexadecane.

Additional file 5: Supplementary Table S5. Presence of W-PE assimilation pathway in metagenomes of wood-, PE-, W-PE-colonising biofilms and the surrounding river water.

## ACKNOWLEDGEMENTS

The authors thank Gabriel Martorell Crespí and Francisca Orvay Pintos from the Scientific-Technical Services of the UIB for the acquisition and analysis of metabolomic and proteomic data.

## AUTHORS’ CONTRIBUTIONS CRediT

**Theo Obrador-Viel**: Conceptualisation, Methodology, Software, Validation, Formal analysis, Investigation, Data curation, Validation Writing – original draft. **Rocío D. I. Molina**: Methodology, Validation, Investigation, Writing – review & editing. **Robyn J. Wright**: Methodology, Software, Formal Analysis, Investigation, Data curation, Writing – review & editing. **Maria del Mar Aguiló-Ferretjans**: Methodology, Investigation, Writing – review & editing. **Vinko Zadjelovic**: Formal Analysis, Resources, Writing – review & editing. **Balbina Nogales**: Conceptualisation, Validation, Writing – review & editing, Supervision, Funding acquisition. **Rafael Bosch**: Validation, Formal Analysis, Supervision, Funding acquisition, Writing – review & editing. **Joseph A. Christie-Oleza**: Conceptualisation, Methodology, Validation, Formal analysis, Resources, Writing – original draft, Visualisation, Supervision, Project Administration, Funding acquisition.

## AVAILABILITY AND DATA MATERIALS

### ETHICS APPROVAL AND CONSENT TO PARTICIPATE

Not applicable.

## CONSENT FOR PUBLICATION

Not applicable.

